# PopTargs: A database for studying population evolutionary genetics of human microRNA target sites

**DOI:** 10.1101/613372

**Authors:** Andrea Hatlen, Mohab Helmy, Antonio Marco

## Abstract

There is an increasing interest in the study of polymorphic variants at gene regulatory motifs, including microRNA target sites. Understanding the effects of selective forces at specific microRNA target sites, together with other factors like expression levels or evolutionary conservation, requires the joint study of multiple datasets. We have compiled information from multiple sources and compare it with predicted microRNA target sites to built a comprehensive database for the study of microRNA targets in human populations. PopTargs is a web-based tool that allows the easy extraction of multiple datasets and the joint analyses of them, including allele frequencies, ancestral status, population differentiation statistics and site conservation. The user can also compare the allele frequency spectrum between two groups of target sites, and conveniently produce plots. The database can be easily expanded as new data becomes available and the raw database as well as code for creating new custom made databases are available for downloading. We also describe a few illustrative examples.

**Availability and implementation:** Poptargs is available at http://poptargs.essex.ac.uk

## INTRODUCTION

As genome sequencing costs continue to decrease, the interest in population genetics increases. In particular, the analysis of variation at regulatory sites is becoming critical to understand how non-coding sequences emerge and evolve (1). MicroRNAs are important gene regulators that target gene transcripts by partial complementarity (2). The fact that their targets can be predicted from their primary sequence has been exploited to study the potential impact of single-nucleotide polymorphisms at their target sites. Indeed, a number of studies have reported selective pressures at these target sites by investigating the variation in populations (3–6).

A number of databases for analyzing polymorphic microRNA target sites exists (e.g. (7)). However, these databases are designed to explore the functional and biomedical implications of single-nucleotide polymorphisms. Despite the interest in population genetics at microRNA target sites, there is currently not a dedicated platform to study evolutionary and population genetics at canonical microRNA target sites. Here we aim to fill this gap. We have developed a database which cross-links allele frequencies and other variables of evolutionary interest at predicted microRNA target sites, as well as expression and evolutionary conservation information from other sources, permitting the analysis of frequency spectrums and population differentiation at target sites.

## METHODS

### Source of data

The human 3’UTRs were downloaded with Biomart (8) and the BiomaRt R package (9) from Ensembl database version 96 (human genome assembly GRCh38), and keeping only 3’UTRs from protein coding transcripts. All mature human microRNAs were downloaded from miRBase version 22 (10). SNPs were also retrieved from the 1,000 Genomes Project (11) as compiled in dbSNP build 151 (12) [Ensembl Variation 96]. Genes were classified as ‘over-’ or ‘under-expressed’ by tissue according to the Bgee database, version 14.0 (13). MicroRNA tissue expression information was obtained from five RNAseq dataset from Meunier et al (2013) and 46 datasets cataloged in MiRmine (15) (accesion numbers are listed in Supplementary Table 1). The microRNA data was classified into four groups for analysis, based on their expression in each tissue: 1- zero RPM (reads per million), 2- broad expression (>50RPM), 3- high expression (>500RPM), and 4- specifically expressed in one tissue (highly expressed compared to the other tissues: 1.5 times the interquartile range plus the upper quartile across tissues. Target and near-target (one nucleotide difference with a target) sites were found using seedVicious 1.1 (16), which predicts canonical target sites without filtering out for sequence conservation. Only SNP locations in which one allele was a target and another allele was a near-target were further considered. This important feature allows the study of target sites that are not in the reference genome, but that can be targets in some populations (see Results and Discussion).

### Access and Implementation

The database is build in MySQL and it is freely accessible via a dedicated web portal at https://poptargs.essex.ac.uk/. The database provides three main options to explore microRNA target sites. First, users can search (*Search* tab) specific microRNAs or genes, or compare the allele frequencies between two lists of microRNAs or genes (*User lists* tab). The web form also gives the option to plot the allele frequencies side to side to a fast visual inspection of results. In the computation of these plots only unique SNPs are used to avoid duplicated results, and p-values from the two one-tailed Kolmogorov-Smirnov tests are provided for convenience. Alternatively, the users may browse the database (*Browse* tab) and select microRNAs with specific expression profiles and/or sequence conservation. This data can be retrieved for all or for specific human populations. The database also provide computations for target sites in the reverse complement strand to the transcript, which can be used as background distributions for statistical purposes. Finally, the user has the option to download the whole MySQL database (*Downloads* tab). Researchers can also create their own databases with custom sequences as we also provide the source code and full instructions at https://github.com/ash8/PopTargs

## RESULTS AND DISCUSSION

The basic search function of PopTargs is the ‘Search’ form. Users can look for microRNAs or genes (Ensembl unique IDs) to find out potential polymorphic sites in which one of the targets is a target site. For each target site the output reports the following features:

- Gene, transcript, and SNP accession numbers, with links to the data source.
- SNP chromosome and position with a link to the UCSC genome browser (17).
- Ancestral and target alleles, together with allele and derive allele frequencies.
- PhyloP scores (average for the whole target site) as pre-computed in UCSC (18).

In addition, for each microRNA the database reports whether it is catalogued as ‘high-quality’ in miRBase, and whether it exists in miRGeneDB (19) and if the mature sequence is the same or not between these two databases. Users can also provide lists of mature microRNAs names and gene names in the *User lists* form.

As we considered near-targets (see above) during the database assembly the user will also find target sites that are not in the reference genome yet one of the alleles is associated to a target site. This feature can be exploited to detect putative target sites not present in the current reference genome sequence (see discussion at the end of this section). The table provides the population frequencies of the target allele, and also reports which allele is ancestral to human populations. Lists of microRNAs of interest can be obtained from miRBase (10) but also from curated databases that may allow the filtering of microRNAs based on evolutionary conservation or other features (e.g. mirGeneDB (19)). The possibility of providing lists of both microRNAs and genes helps to narrow down the targets of interest when a specific subset of experimentally validated interactions (for instance, from TarBase (20) or miRTarBase (21)) is to be explored. The database also allows the possibility of plotting allele frequencies for the queried microRNA/gene interactions. In this case, one can plot the allele frequencies at target sites and compare it with the allele frequencies of either an alternative list of microRNAs or an alternative list of genes. This is particularly handy when visually exploring large amount of data (see below).

To explore variation at target sites in pre-computed lists, the ‘Browse’ form allows to study microRNAs with different levels of expression, expression breath, evolutionary conservation and even sub-population structure. For instance, we recently reported that in human populations there is detectable selection against microRNA target sites (6). We can explore some specific cases with PopTargs. If we use the *Browse* option we can compare target sites for microRNAs highly expressed in testis (for instance) versus microRNAs not detected in testis. PopTargs will produce an allele frequency and a derived allele frequency plots, showing that the frequency of the target allele is significantly lower for the targets of highly expressed microRNAs (Figure 1). This result suggests that when a target site for a testis microRNA randomly appears in a testis expressed gene, there will be selective pressures to remove this allele from the population.

**Figure 1.**
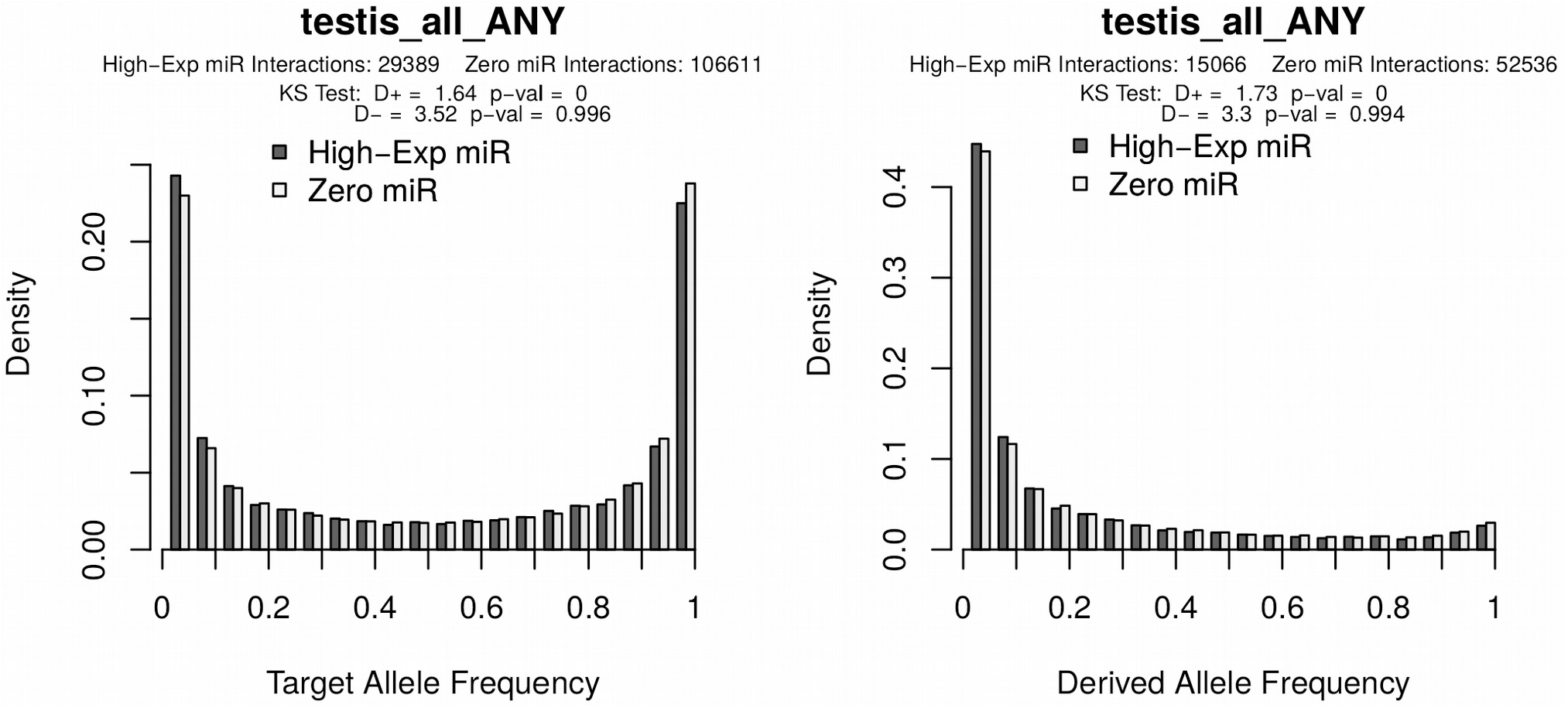
Allele frequency distributions as generated from the PopTargs web server. The left panel show the target allele frequency distribution for microRNAs highly expressed in testes (grey bars) and for microRNAs whose expression was not detected in testes (white bars). Likewise, the right panel shows the target allele frequency distribution of derived alleles, that is, where the ancestral allele is a non-target. The latter plot is also often called the Site Frequency Spectrum.

We can download a full table with the results, which will contain allele and derive allele frequencies, but also the target allele frequencies for different human populations and the estimated Fst (22). From the results produced we can detect unique 12 segregating target:non-target allele pairs for microRNAs highly expressed in testis (Table 1), that have a high degree of population differentiation (Fst>0.7). For instance, transcripts from *MTAP* gene has a conserved target site for let-7a-5p, but this target site is not detected in the reference genome. Indeed, the loss of the ancestral target site happened in European populations whilst other human groups mostly maintain the target allele (dbSNP entry rs6912739, Table 1). This result illustrates how population dynamics can be used to detect target sites that are not in the reference genome and, therefore, escape most target prediction programs (Helmy, Hatlen, Marco, under review).

**Table 1:** Target sites for testis-expressed microRNAs with a high degree of population differentiation.

| **MicroRNA** | **Gene** | **SNP** | **target is**<br>**ancestral** | **EAS** | **AMR** | **AFR** | **EUR** | **SAS** | **all** | **Fst** |
| --- | --- | --- | --- | --- | --- | --- | --- | --- | --- | --- |
| miR-202-5p | ATP1A1 | rs1885802 | yes | 0.0427 | 0.1268 | 0.8109 | 0.0338 | 0.0450 | 0.2559 | 0.7914 |
| miR-130a-3p | SLC30A9 | rs12511999 | yes | 0.0486 | 0.2363 | 0.9130 | 0.2495 | 0.2219 | 0.3773 | 0.7302 |
| miR-513a | TCERG1 | rs3822506 | no | 0.7183 | 0.1167 | 0.0325 | 0.0915 | 0.2198 | 0.2307 | 0.7224 |
| miR-151a-3p | MTAP | rs12003714 | no | 0.9921 | 0.9380 | 0.2042 | 0.9950 | 0.8824 | 0.7567 | 0.8638 |
| let-7a-5p | MTAP | rs7875199 | yes | 0.0536 | 0.0576 | 0.7958 | 0.0070 | 0.1288 | 0.2555 | 0.7997 |
| miR-24-3p | SCN2B | rs624328 | no | 0.9405 | 0.8905 | 0.2519 | 0.9513 | 0.9673 | 0.7601 | 0.7674 |
| miR-192-5p | C12orf65 | rs1533703 | yes | 0.9980 | 0.7262 | 0.1490 | 0.7694 | 0.7945 | 0.6512 | 0.7074 |
| miR-25/92a-3p | PTK6 | rs186332 | no | 0.9643 | 0.8732 | 0.0469 | 0.9284 | 0.5930 | 0.6304 | 0.8487 |

The target allele frequencies are provided for East Asian (EAS), Mixed American (AMR), African (AFR), European (EUR) and South Asian (SAS) populations, as described in the 1000 genomes project (see Methods).

We provided all scripts used to generate the original database and full documentation such that interested users can generate their own database. As the number of available genome sequences increases, this feature can be of use to those interested in expanding the current database.

## ACKNOWLEDGMENTS

We thank Stuart Newman for his invaluable help in setting up the server to host our database and web portal.

## FUNDING

This work was supported by the Wellcome Trust [grant number 200585/Z/16/Z].

## Notes

#### Summary of Updates

New version submitted for revision. Also, it describes the new version 2 of the database with expanded datasets.

https://poptargs.essex.ac.uk/

