## Supplementary Table 1 for "PopTargs: A database for studying population evolutionary genetics of human microRNA target sites"

**Supplementary Table 1.** Accession number of sRNA-seq experiments processed in this study.

| Tissue | EBI-ENA accession number |
|---|---|
| Blood | DRX012360; DRX012361; DRX012362; SRX318164; SRX318180; SRX386679; SRX386680; SRX386681; SRX426123; SRX426124; SRX426125; SRX426475; SRX426478; SRX666575; SRX666576; SRX666577; SRX666578; SRX666579 |
| Brain | SRX182778; SRX375448; SRX375450; SRX375452; SRX375455; SRX375461; SRX375462; SRX375463; SRX375464; SRX375466; SRX375467 |
| Breast | SRX513283; SRX513284; SRX513285; SRX513286 |
| Cerebellum | SRX182779 |
| Heart | SRX182780 |
| Kidney | SRX182781 |
| Liver | SRX353113 |
| Lung | DRX003170; DRX003171 |
| Placenta | SRX262196; SRX262197; SRX262198; SRX262199; SRX262200; SRX262201; SRX262202; SRX262203 |
| Testis | SRX182782; SRX271415; SRX271416; SRX271417 |
